# New approaches to the Single-Interval Adjustment Matrix (SIAM) Yes-No task

**DOI:** 10.1101/2024.02.27.582322

**Authors:** Daniel Shepherd, Michael J. Hautus

## Abstract

Two adaptations of the Single-Interval Adjust-Matrix Yes-No (SIAM-YN) task, designed to increase the efficiency of absolute threshold estimation, are described. The first, the SIAM Twin Track (SIAM-TT) task, consists of two interleaved tracks of the standard SIAM-YN that are run in the same trial with a single response. The second new task modifies the binary SIAM-YN task by using a six-point rating-scale (SIAM-Rating). In Experiment 1, data from three tasks estimating absolute thresholds were obtained using a 10-ms tone, the 2-IFC transformed up-down procedure, SIAM-YN task, and the SIAM-TT task. The data support the use of the SIAM-TT as an alternative to the conventional two-interval and one-interval (SIAM-YN) tasks when used to estimate absolute thresholds. By presenting two interleaved SIAM-YN tracks on a single experimental trial, the SIAM-TT task possesses greater efficiency alongside its signal-detection heritage which confers less response bias. Similarly, in Experiment 2, which compared the 2-IFC adaptive, SIAM-YN, and SIAM-Rating tasks, there was no main effect of task upon threshold estimates. The findings replicate previous studies supporting the validity and efficiency of the SIAM-YN task, and extends the SIAM-YN toolbox to efficiently facilitate the generation of psychometric functions (the SIAM-TT task) and Receiver Operating Characteristic Curves (the SIAM-Rating task).

## Introduction

A psychophysical measure commonly utilised by experimental psychologists is the absolute threshold, which represents in physical units the smallest amount of a stimulus required to be detected according to some operationally defined performance criterion (e.g., percentage correct). There are numerous methods affording threshold estimation, divided into approaches that employ a predefined range of stimuli (e.g., Method of Limits) or those that adjust the stimuli on a trial-by-trial basis (i.e., adaptive methods). In regards the latter, the two-down one-up and three-down one-up staircase procedures [1] are popular due to their greater efficiency when compared to approaches using fixed-level stimulus sets. In the detection context, these staircase procedures typically involve a two-interval forced-choice (2-IFC) structure, whereby a single trial presents two temporally distinct observation intervals in which one has been randomly assigned the target stimulus. The 2-IFC adaptive task consists of a series of trials in which, for all but the first trial, the intensity of the stimulus is adjusted according to the participant’s response history. The same is true for the Single-Interval Adjustment Matrix Yes-No (SIAM-YN) task [2], though in the literature the SIAM-YN task does not enjoy the same popularity as its 2-IFC counterparts.

In the hearing literature, 2-IFC adaptive procedures are more widely reported than single-interval procedures. However, some have argued that single-interval methods are higher in statistical efficiency and lower in statistical bias [3,4,5,6,7]. Two-interval tasks, for example, obtain the same amount of information as single-interval tasks, but with the penalty of an extra observation interval. In the detection context, Treutwein [8] argued for tasks capable of producing valid and reliable threshold estimates while being intuitive to the participant. However, depending on the task or stimuli, valid estimates of threshold can require large amounts of data in order to minimise measurement error, though excessive trials can lead to fatigue or are not always feasible for practical reasons [9]. The SIAM-YN task is a relatively new single-interval detection task developed by Kaernbach [2], which has been independently validated [9, 10] and modified [10]. It is a single-interval task requiring the participant to indicate whether a target stimulus was presented during a trial using a binary yes-or-no response.

The theoretical underpinnings of the SIAM-YN task are elucidated by its creator [2] and elsewhere [9, 10], and only a cursory description will be offered here. At the centre of the SIAM-YN task is the adjustment matrix, which can be considered in the same light as a traditional pay-off matrix. However, rather than consisting of some tangible (e.g., food or money) reinforcer or punisher, the adjustment matrix rewards or punishes through the adjustment of signal intensity. Thus, a key assumption of the SIAM-YN task is that, because the participant is motivated to maximise their performance, non-biased (i.e., neutral response criteria) performance can be achieved via trial-by-trial feedback. The 2 x 2 adjustment matrix, in turn, is determined by the target performance (*t*) set by the investigator, where *t* represents the maximum distance between the Receiver Operating Characteristic (ROC) curve and the major diagonal. This distance (also called the *maximum reduced hit rate*) must, for a symmetrical ROC, fall on the minor diagonal, where the slope of the ROC curve equals one. Hence the SIAM-YN procedure involves “chaperoning” a participant’s operating point in ROC space to where the slope of the ROC representing an unbiased observer, while simultaneously adjusting signal intensity to match *t*.

The current study extends the SIAM-YN task in two ways and evaluates these modifications. Firstly, in order to reduce the time taken to estimate absolute thresholds we modify the SIAM-YN task by doubling the number of observation intervals and presenting two independent SIAM-YN tracks in a single trial. We denote this task the SIAM-TT (twin track) to differentiate this task from the orthodox SIAM-YN task. As each response at the end of a trial contributes to the estimation of two threshold measures the reduction in warning, response, and feedback intervals can potentially provide large gains in efficiency over conventional single or two-interval tasks. Secondly, to avoid the constraints and short-comings of binary-response regimes, we converted the SIAM-YN task to a rating-scale task with six response categories, denoted the SIAM-Rating task. In the SIAM context, this adaptation allows the consequences of trial-by-trial decisions to be weighted; that is, more rewarding or more punishing.

While adaptive procedures originating from psychophysics are utilised within the width and breadth of experimental psychology, little innovation has occurred in the area in the last few decades, with the SIAM-YN task itself now over thirty years old. To assess the effectiveness of the SIAM-TT and the SIAM-Rating tasks, two experiments were performed that involved the collection of human data. Both experiments involved procuring data from the two-interval forced-choice (2-IFC) transformed up-down procedure and the standard SIAM-YN task, with the former task being treated as a gold standard to which other adaptive procedures can be benchmarked [2, 11, 12, 13]. Lastly, this study offers a further opportunity to validate the SIAM-YN task in the auditory context.

### Experiment 1: The SIAM-TT Task

The SIAM-TT task is similar to the orthodox SIAM-YN task, but has two key procedural differences. First, whereas the SIAM-YN task pauses while waiting for the participant to respond, the SIAM-TT does not. Instead, the SIAM-TT mimics the go/no go task and incorporates aspects of the Method of Free Response [14]. Practically speaking, the SIAM-TT task does not have an indefinite response interval, and instead extracts information from both a response (i.e., a button press) and non-response (i.e., no action) to adjust the stimulus magnitude for the next trial. The Method of Free Response has successfully been incorporated into the SIAM-YN task previously, notably the single-interval SIAM-Rapid task [10]. Second, the SIAM-TT task has two observation intervals, however, on any one trial the task of the participant is to indicate if a stimulus occurred in only one, or both, or neither, of the two intervals. Further, across a block of SIAM-TT trials there are two interleaved SIAM-YN tracks, one ascending and one descending, and on any one trial the tracks are assigned randomly to either of the two observation intervals. The inclusion of both ascending (i.e., track begins with a subthreshold stimulus) and descending (track begins with a suprathreshold stimulus) is justified on two grounds. Firstly, to avoid potential loss of independence between the two tracks through high covariance. Pertinently, if both tracks start at the same stimulus level the participants may use information from one track to inform decision-making on the other, thus biasing responses. Secondly, this approach allows the generation of a full psychometric function.

For the purpose of evaluating the SIAM-TT task, psychometric functions will be constructed from which a further estimate of threshold can be derived. A disadvantage of adaptive methods is that they do not produce a full psychometric function, usually because stimulus values displaced from the threshold are represented by only a few experimental trials. This issue is compounded by the usual practice of starting adaptive tracks with suprathreshold stimuli, rather than subthreshold stimuli. A solution is to have both ascending (subthreshold starting level) and descending (suprathreshold starting level) tracks. While interleaving multiple adaptive tracks is in itself not novel, the tracks are typically isolated and presented in a one-track-per-trial fashion. By presenting two-tracks-per-trial, the SIAM-TT task retains the advantages of using multiple tracks (e.g., reducing predictability and aiding memory) while avoiding the primary disadvantage: requiring twice as many trials to obtain two independent estimates of threshold.

## Method

Twenty seven inexperienced participants, 11 males (*M*_*age*_ = 24.18, *SD* = 3.40) and 16 females (*M*_*age*_ = 27.31, *SD* = 9.49) participated in the study. Potential participants were excluded if they reported current or historical hearing pathology, or other major health problems. This study along with Experiment 2 was approved by **<< Blinded for Review >>** Human Participants Ethics Committee. A repeated-measures design was adopted involving three types of detection task: the SIAM-TT task and two benchmarking tasks: the 2-IFC 3-down 1-up adaptive procedure and the SIAM-YN task. Here, we seek to determine if statistically equivalent estimates of absolute threshold can be obtained across the three tasks. In total, participants underwent 30 blocks of trials, ten blocks for each task, with the tasks randomly presented across the experimental series.

The stimulus to be detected was a 1000-Hz tone of 10-ms duration with 1-ms ramps (cos^2^). Tones were generated digitally using LabVIEW 8.1 (National Instruments) and converted to analogue using a sampling rate of 44.1 kHz. The sound pressure level of the tone was controlled by a programmable attenuator (Tucker Davis Technologies, TDT, PA5) which then routed the tone to a monaural earpiece (Telephonics, TDH-49P) via a headphone buffer (TDT HB7). All participants received the stimuli in the left ear. For all three tasks a descending track commenced with a 40 dB SPL suprathreshold tone, while for the SIAM-TT ascending track the starting level was 5 dB SPL. These starting levels were determined by a pilot study using a single naïve participant yielding an absolute threshold of approximately 23.5 dB SPL.

Participants were seated in a sound-attenuating chamber (Amplaid, Model E) in front of a set of light emitting diodes (LEDs) functioning as warning and feedback lights. Responses were made using a custom-built button box. Prior to beginning the experiment the participants were briefed on the three types of task (i.e., 2-IFC, SIAM-YN, and SIAM-TT tasks) and provided laminated instruction sheets which they kept with them when undertaking each task. For all three tasks trial-by-trial feedback was provided, indicating either correct or incorrect responses.

For the 2-IFC adaptive procedure each trial consisted of two observation intervals, with each having an equal chance (*p* = .5) of being assigned the tone. A trial began with a 400-ms warning light and then a 400-ms pause before the first and second observation intervals, punctuated by a 400-ms inter-stimulus interval, were presented. The observation intervals were 10 ms in duration and marked by the illumination of a green LED. In the ensuing response interval participants were required to use the button box to indicate which of the two intervals contained the tone, with feedback provided by the LEDs contingent on response. In accordance with the 3-down 1-up task, an incorrect response increased the level of the tone by 1 dB, while three consecutive correct responses decreased the level of the tone by 1 dB. A block of trials terminated after 15 turnarounds, with the absolute threshold taken as the average of the last 12 turnarounds. A turnaround occurs when the sequence of stimuli reverse from an ascending to a descending series of stimulus levels, or vice versa [15].

For the SIAM-YN task a single observation interval, marked by a 400-ms LED, was presented per trial. On any one trial there was a 50 percent chance that the tone would be present. Each trial began with a 400-ms warning LED and 400-ms pause, followed by a 10-ms observation interval also marked by an LED. During the response interval the participant indicated if the tone was present using the button box, after which feedback was provided and the next trial began. As per the stipulates of the SIAM matrix (*t* = .5: [2]), a Hit reduced the level of the tone by 1 dB, while a False Alarm or a Miss increased it by 2 dB and 1 dB, respectively. A Correct Rejection left the level of the tone unchanged. As with the 2-IFC adaptive task described above, a block of trials terminated after the 15^th^ turnaround.

The SIAM-TT task possesses two observation intervals, each containing an independent SIAM-YN track. On any one SIAM-TT trial these interleaved tracks are randomly assigned to either the first or second observation interval. A SIAM-TT trial began with a 400-ms warning LED, and then two 50-ms observation intervals, separated by a 400-ms inter-stimulus interval. This addition of an extra observation interval included in a SIAM-TT trial demands a modified response interval. Whereas the interval duration for the response interval for the SIAM-YN task is determined by the participant (i.e., the next trial is contingent upon response), the SIAM-TT has a fixed duration response interval of three seconds. During this time, participants indicate if a tone was sensed in both intervals (simultaneous press of left and right buttons), in the first (left button) or second (right button) intervals only, or if no tones were perceived to be present during the trial (no buttons pressed). For the purposes of the current study, 106 SIAM-TT trials per block were obtained, so as to facilitate the construction of psychometric functions. However, absolute thresholds were estimated as per the 2-IFC adaptive and SIAM-YN tasks, thus calculated by taking the average of the 4^th^ to the 15^th^ turnarounds. Figure 1 plots data from a single SIAM-TT block performed by one of the authors.

**Fig. 1.**
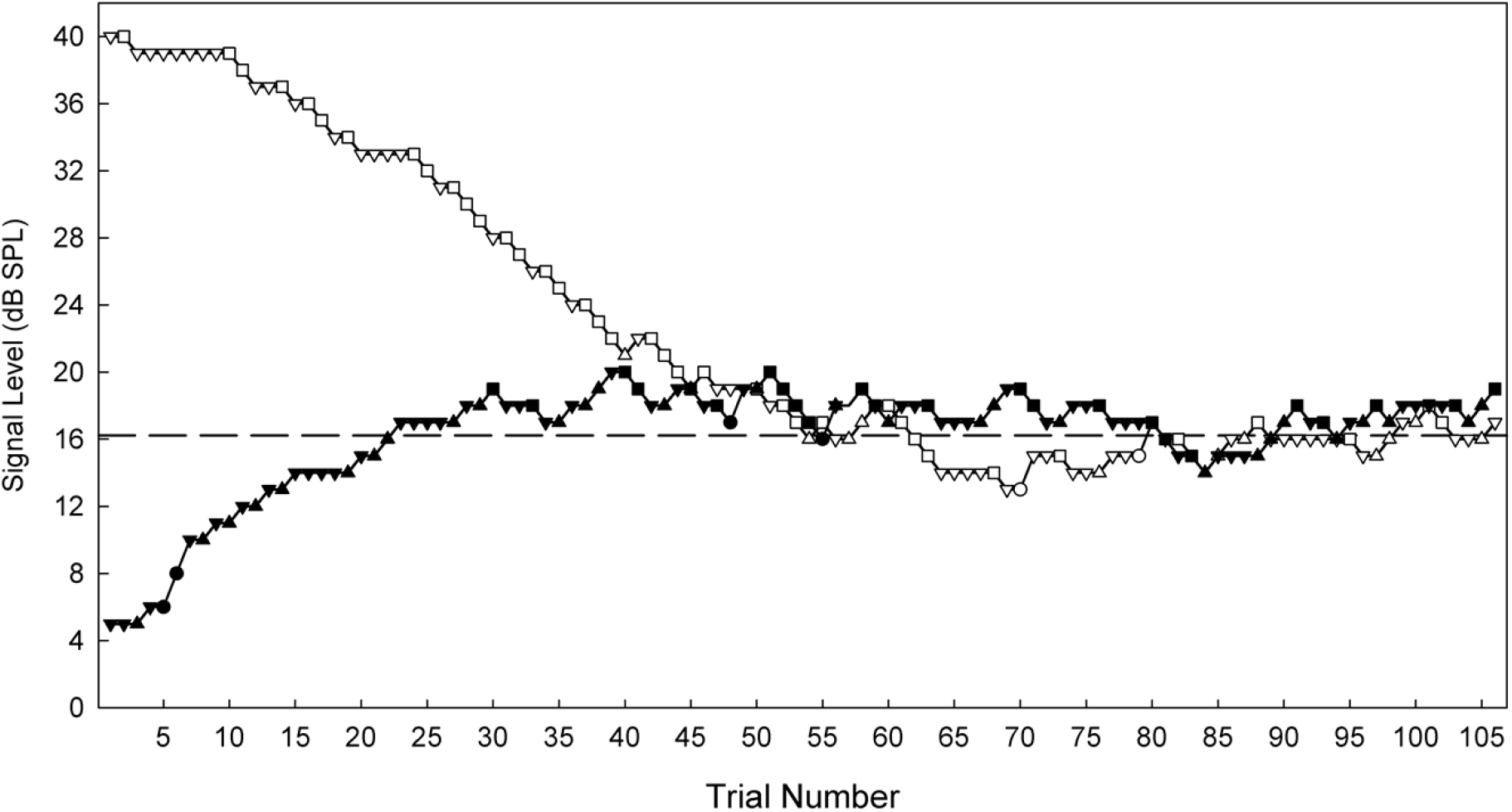
Signal level as a function of trial number for a single SIAM-TT block obtained by the first author during testing. The ascending track is represented by filled symbols, and the descending track by open symbols. For both tracks the squares (▪) represent hits, triangles misses (▲), circles false alarms (●) and inverted triangles (▼) correct rejections. The dashed horizontal line is the threshold (16.21 dB SPL) calculated for this block of trials. Note that the descending track (open symbols) would mirror a track from the conventional SIAM-YN task.

## Results

### Absolute Thresholds

Table 1 displays mean (*M*) thresholds and associated standard deviations (*SD*) for the three tasks, with the SIAM-TT estimate being calculated as the mean of the ascending and descending thresholds. These values of approximately 20 dB SPL compare favourably to those reported in the literature using the same stimuli [16]. A factorial repeated-measures ANOVA using Task (three levels) and Block (ten levels) determined that there were no significant differences in mean threshold estimates across the three tasks (*F*(2, 52) = 1.475, *p* = .238, η ^p2^ = .054), indicating good convergent validity. Figure 2 indicates the degree of convergent validity across the three tasks, which is slightly stronger than that reported previously [10] for the 2-IFC and SIAM-YN tasks. Block was, however, significant (*F*(9, 234) = 7.665, *p* < .001, η _p_^2^ = .228), and is explained by the anticipated learning effects, which were equivalent across Task as evidenced by the lack of a significant interaction effect (*F*(18, 468) = 1.419, *p* = .117, η_p_^2^ = .052).

**Table 1.** Mean absolute thresholds (dB SPL) across the three detection tasks of Experiment 1.

| Task | Min | Max | Range | Mean | SD |
| --- | --- | --- | --- | --- | --- |
| 2-IFC Task | 15.48 | 26.57 | 11.09 | 20.40 | 2.185 |
| SIAM-YN Task | 15.93 | 26.51 | 10.58 | 20.77 | 1.774 |
| SIAM-TT Task | 15.77 | 27.22 | 11.45 | 20.64 | 1.611 |

**Fig. 2.**
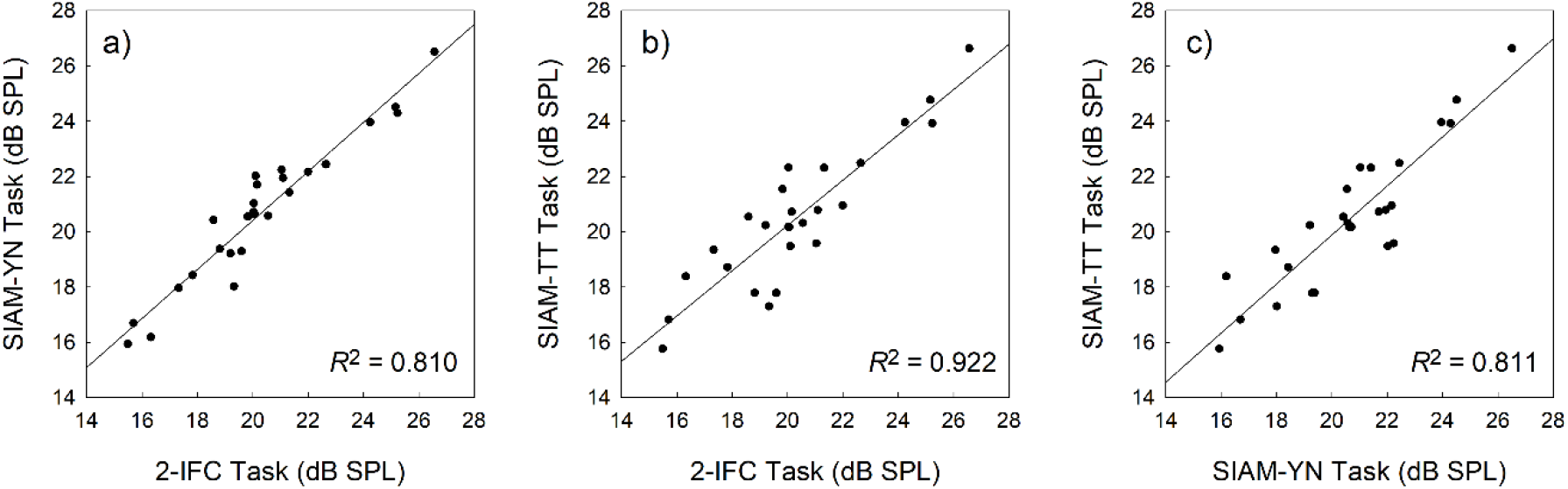
Scatterplots exhibiting the association between thresholds obtained with the 2-IFC and SIAM-YN tasks (a), the 2-IFC and SIAM-TT tasks (b), and the SIAM-YN and SIAM-TT tasks (c). The solid lines represent the best linear least-squares fits, and the accompanying coefficients of determination suggest collinearity.

Comparisons of the standard deviations across the three tasks suggests that the SIAM-TT threshold estimate is as reliable as those obtained with the 2-IFC and SIAM-YN tasks. It is argued that in the hearing-threshold context an inverse relationship between standard deviation and reliability exists [17], and Kaernbach [2] reported SIAM-YN thresholds to be less variable than those from 2-IFC adaptive procedures, a finding that is replicated here. A repeated-measures ANOVA showed significant differences (*F*(2,26) = 6.386, *p* = .018, η _p_^2^ = .161) across Task in terms of mean standard deviations, with *post hoc* tests indicating that the 2-IFC standard deviation was significantly greater than that for the SIAM-YN task (*p* = .041), but not the SIAM-TT task (*p* = .054).

### Psychometric Function

A conventional, non-adaptive, representation of the SIAM-TT data can also be generated in terms of the psychometric function, which plots the relationship between proportion correct and stimulus level. For each participant, data were pooled across the ten SIAM-TT blocks and used to generate empirical psychometric functions, to which theoretical functions of the form

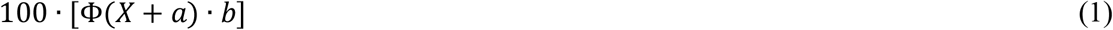

were then applied. Here, Ф represents the cumulative standard normal distribution, and the best-fitting parameter estimates of *a* and *b* were obtained using maximum likelihood estimation. The parameter *a* permits the psychometric function to move laterally while *b*, determining the slope of the function, is necessary because the 50% point on the function can shift with slope. Figure 3 shows data pooled across the entire sample, where the area of a data point represents the number of trials on which that point is based (*M* = 741.74, *SD* = 758, *Min* = 30, *Max* = 2407). The goodness-of-fit for the function in Figure 3 was *R*^2^ = 0.99, while the mean goodness-of-fit across the 27 individual functions was *R*^2^ = 0.96 (*SD* = 0.04). Adopting the standard performance criterion for the single-interval yes-no task, 50% correct detections, a value of 19.82 dB SPL can be calculated from the best-fitting curve in Figure 3. This value is within 1 dB of those reported in Table 1, and a further repeated-measures ANOVA performed on individual data indicated no statistically significance differences in thresholds derived from Equation 1 and those obtained from the 2-IFC and SIAM-YN tasks (*p* > .05).

**Fig. 3.**
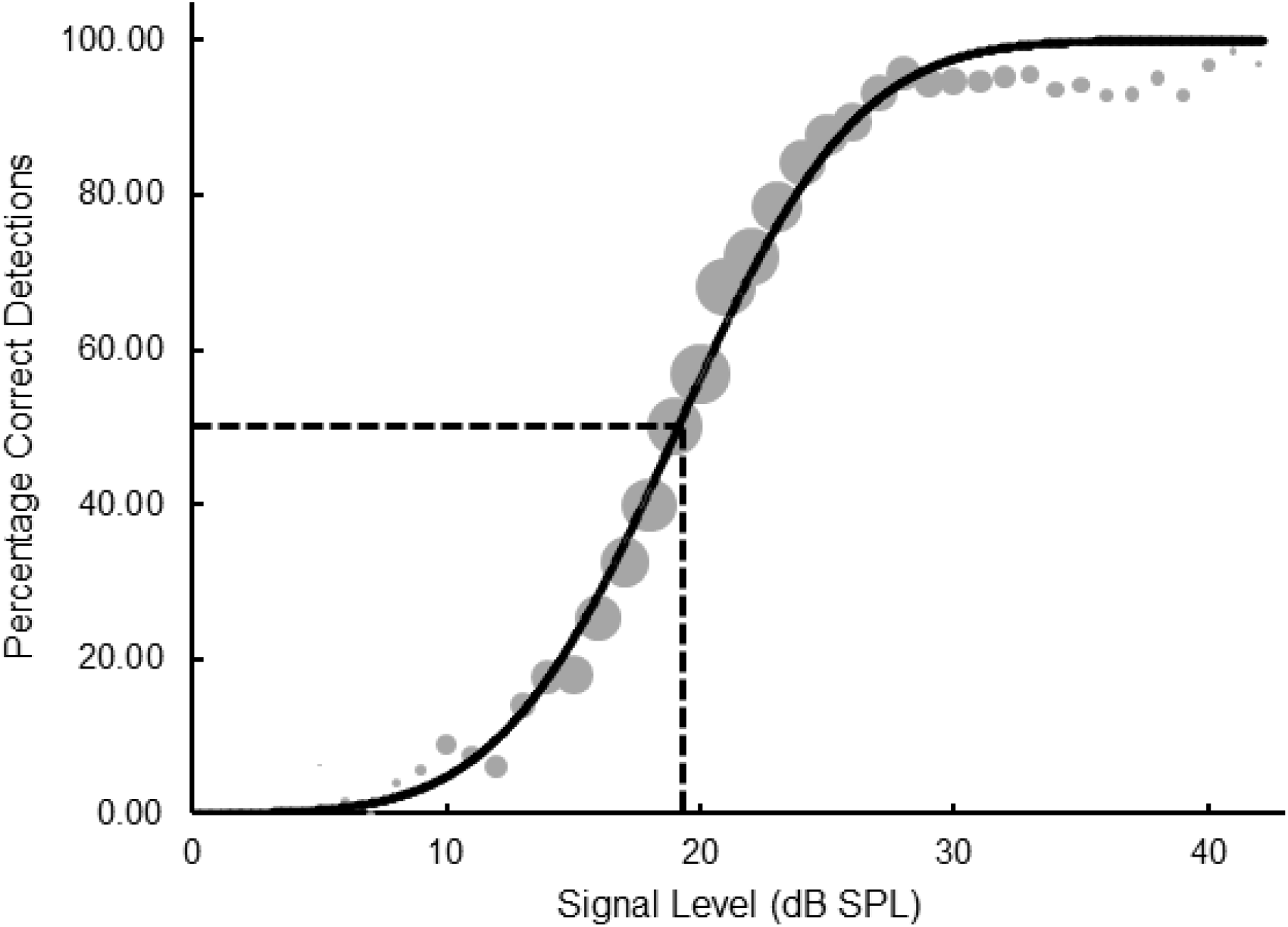
Psychometric function plotting percentage correct detections as a function of signal level for pooled data obtained from the SIAM-TT task. The slope parameter is equal to 1.18.

### Temporal Analysis

Table 2 presents the mean number of trials and mean time (seconds) it took each task to achieve 15 turnarounds. The accuracy of these figures is slightly biased, as the software only time-stamped the beginning of a trial to the nearest second. Additionally, because the ascending and descending tracks in the SIAM-TT approached 15 turnarounds at different rates, the data for each track is analysed separately. The superscripts in Table 2 indicate, across a single row, significant differences across the three tasks. Of note, the 2-IFC task required a significantly greater number of trials than the SIAM-TT task, though not the SIAM-YN task. Regarding seconds-per-trial, the SIAM-TT was, as anticipated, significantly longer than the 2-IFC and SIAM-YN tasks. However, the important point to remember is that the SIAM-TT is returning two, as opposed to one, threshold estimate, for only a small investment of more time (i.e., 4.2 seconds/trial). As an approximation, it would take the SIAM-YN task an average of five (2.53 x 2) seconds to complete two trials, while for the 2-IFC task this would be approximately six seconds. The differences in total trials between the ascending and descending SIAM-TT tracks is explained by the descending track having a starting point 20 dB SPL greater than the mean SIAM-TT threshold estimate, while the ascending track was only 15 dB SPL below. Finally, it was interesting to note that the SIAM-TT measures had significantly lower trials/turnaround ratios that either of the 2-IFC or SIAM-YN task. While this may be due to some yet investigated factor such as task difficulty, the threshold estimates associated with the SIAM-TT were none-the-less statistically equivalent to those of the 2-IFC and SIAM-YN tasks.

**Table 2.**
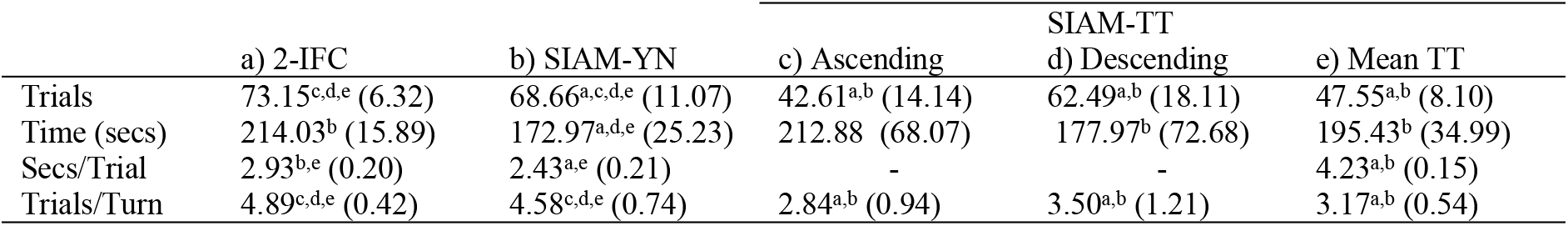
Group means indicating the number of trials, time taken, seconds-per-trial, and trials-per-turnaround, across the three tasks. Superscripts should be referenced across a single row, and indicate Bonferroni-adjusted significant differences across tasks (*p* < .017).

**Table 3.** Mean absolute thresholds (dB SPL) calculated for pooled data across the three detection tasks performed as part of Experiment 2.

| Task | Min | Max | Range | Mean | SD |
| --- | --- | --- | --- | --- | --- |
| 2-IFC Task | 14.58 | 29.45 | 14.87 | 19.97 | 1.723 |
| SIAM-YN Task | 16.32 | 29.35 | 13.03 | 20.20 | 1.891 |
| SIAM-Rating Task | 15.49 | 27.63 | 12.14 | 19.75 | 1.822 |

### Experiment 2: The SIAM-Rating Task

In a confidence-rating task the participant is asked to rate their confidence that, assuming a single-interval task, a target stimulus had been present. Using this method with a standard Yes-No task an entire Receiver Operating Characteristic (ROC) curve can be generated. Note that, within reason, as the number of rating categories increases so too does the ability of the experimenter to determine if the selected model is correct for the data. An ill-fitting model can either overestimate or underestimate true sensitivity, so an added convenience of the rating method is that it allows a more thorough test of the theoretical ROC against the data it is supposed to fit. The signal detection index *d*′ can be extracted from the ROC and be converted to the equivalent percentage correct that would be obtained by an unbiased participant from a 2-IFC task using [18]

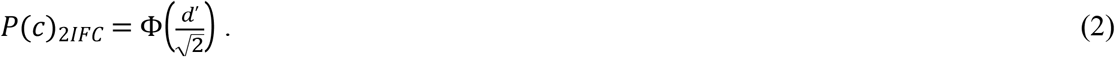

Confidence-rating tasks are typically used with two interval tasks, and appear to have found scant application in single-interval tasks, even though sensitivity measures derived from a confidence-rating task equate to those derived from the Yes-No Task [19]. There has been little recent development in the psychophysical methods used to estimate absolute thresholds, and in the literature the 2-IFC adaptive procedures are more widely reported than single-interval procedures. However, some have argued that single-interval methods are higher in statistical efficiency and lower in statistical bias, or lament the constraint of equal up-and-down stepsizes typical of adaptive procedures [2]. The combination of a confidence-rating response regime and SIAM-YN provides an opportunity to present a detection task with variable stepsizes contingent upon participant response. Figure 4 illustrates the binary nature of the SIAM-YN task (i.e., “Target” vs. “Blank”) and its translation to a confidence-rating regime, included are response outcomes: Hit, Miss, False Alarm (FA), and Correct Rejection (CR).

**Fig. 4.**
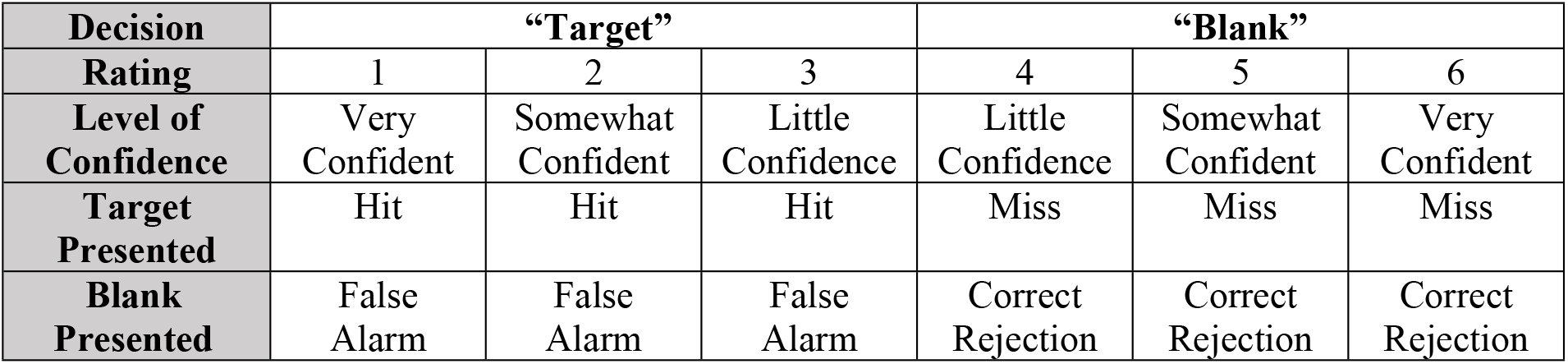
Matrix showing the ratings, their interpretation, and outcome when either the target or the blank is presented.

Additionally, the use of a rating regime permits reckless responses to incur greater punishment if unsuccessful as opposed to cautious responding when the participant is less confident. These outcomes are typically represented in signal detection theory as a payoff matrix. Figure 5 presents the scaling of signal intensity contingent on response. For example, stating that you were very confident that the target was presented when in fact it was not increases the signal intensity by 6 dB, whereas if you were very confident to the contrary then the signal intensity would remain unchanged. The central tenet of the SIAM Yes-No task is that a payoff matrix can be substituted with an adjustment matrix, which adjusts the stimulus to induce the participant to adopt a neutral response criterion via their reinforcement history. The inclusion of a six-point rating-scale allows the adjustment matrix more flexibility with outcomes, and to better match the punishment to the crime or the prize to the victory.

**Fig. 5.**
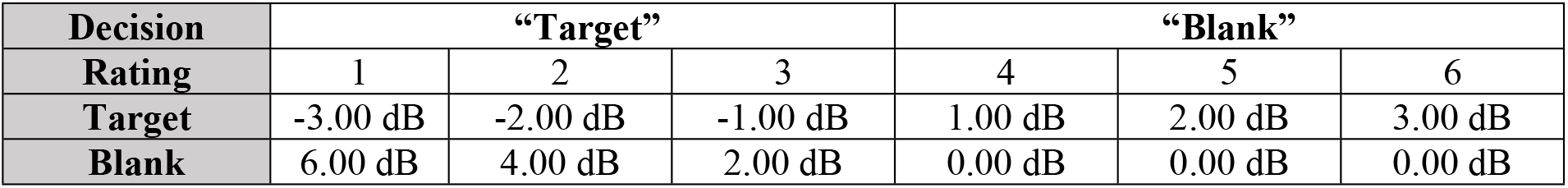
Matrix showing response outcomes expressed in physical units.

### Method

Experiment 2 involved twelve naïve participants, none of whom participated in Experiment 1. There were five males and seven females between the ages of 21 and 25, with none reporting any health issues that might affect their performance. Consistent with Experiment 1 a repeat measures approach was adopted, again employing both the 2-IFC adaptive task and the SIAM-YN task as benchmarks. As such, Experiment 2 was similar to Experiment 1 but with the following differences:

a)instead of assessing the SIAM-TT, Experiment 2 assessed the SIAM-Rating task;

b)as informed by Experiment 1, the starting level on the 2-IFC and SIAM-YN tasks was set to 35 dB SPL;

c)to afford further comparison with Experiment 1, the SIAM-YN task included an even mix of ascending and descending series of trials, allowing the generation of a psychometric function.

Of note, the SIAM-Rating task is identical to the SIAM-YN task used in Experiment 1 in all aspects apart from a modified response interval and associated adjustment. Whereas the SIAM-YN task has a response regime offering the participant a binary choice (i.e., ‘Yes’ or ‘No’), the SIAM-Rating task presents a six-point rating-scale (*re*: Figures 4 and 5).

## Results

### Absolute Thresholds

Mean (*M*) thresholds and associated standard deviations (*SD*) for the three tasks are displayed in Table 3, and as for Experiment 1 are comparable to those reported in the literature [16]. Of note, the threshold estimates for the 2-IFC task and the SIAM-YN task across Experiment 1 (*re*: Table 1) and Experiment 2 (*re*: Table 3) appear comparable. Similarity was determined using independent samples *t*-tests, which returned non-significant probability values for both the 2-IFC (*t*(35) = 0.488, *p* = .629) and SIAM-YN (*t*(35) = 0.679, *p* = 0.501) tasks. Turning back to Experiment 2, differences in mean thresholds across the three tasks were again tested using a 3 (task) x 10 (block) factorial repeated-measures ANOVA. Convergent validity was confirmed by the absence of a significant main effect of task (*F*(2, 22) = 0.982, *p* = .930, η _p_^2^ = .082), and can be assessed graphically in Figure 6. Unlike Experiment 1, there was no significant main effect of block for Experiment 2 (*F*(9, 99) = 1.627, *p* < .118, η _p_^2^ = .129), nor a task x block interaction (*F*(18, 468) = 0.849, *p* = .641, η _p_^2^ = .072). Also different to Experiment 1, a repeated measures ANOVA revealed no significant differences in standard deviation across the three tasks (*F*(1, 11) = 0.239, *p* = .789, η _p_^2^ = .021). These non-significant results this may due to Experiment 2’s smaller sample size, and hence statistic power, of Experiment 2.

**Fig. 6.**
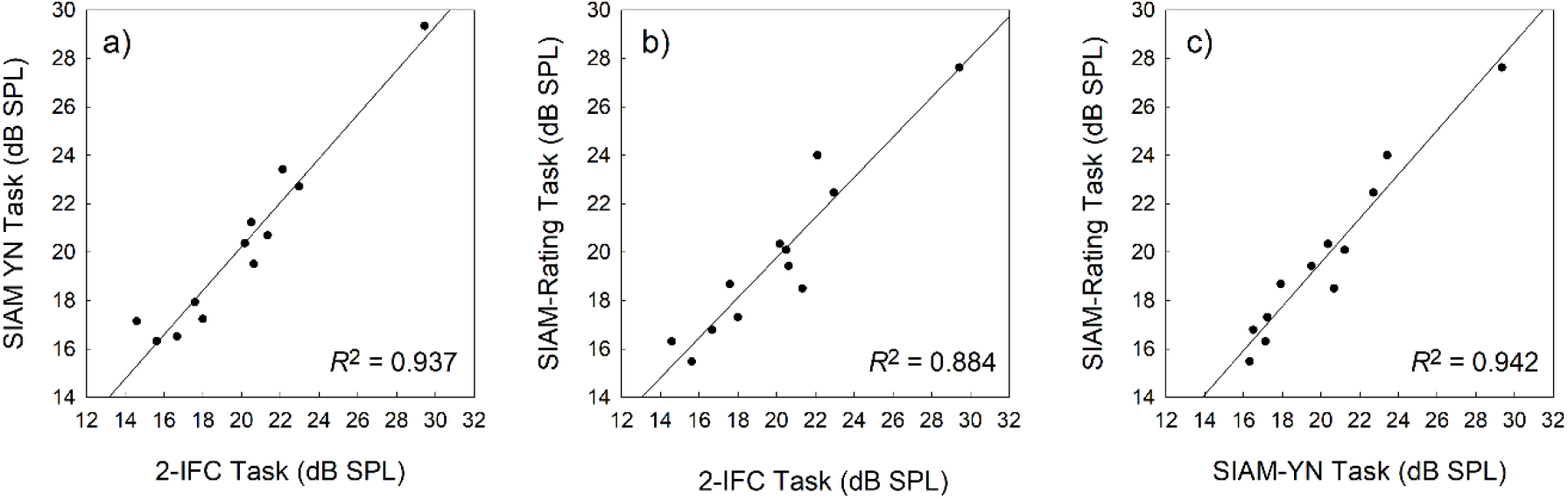
Scatterplots exhibiting the association between thresholds obtained with the 2-IFC and SIAM-YN tasks (a), the 2-IFC and SIAM-Rating tasks (b), and the SIAM-YN and SIAM-Rating tasks (c). The solid lines represent the best linear least-squares fits.

### Receiver Operating Characteristic Analysis

The rating data was analysed using SDT Assistant V1.0 [20], in which criterion-free indices of detection were calculated using maximum likelihood estimation for signal levels between 5 and 35 dB SPL. At each signal level both the normal-normal equal-variance (NN-EV) and normal-normal unequal-variance (NN-UV) models were fitted, thus yielding two estimates of sensitivity per signal level: *d′* and *d*_*a*_ respectively. Examples of the NN-EV and NN-UV models are displayed in Figure 7, where the area under the Receiver Operating Characteristic (ROC) curve is directly proportional to *d′* and *d*_*a*_, representing the participant’s ability to detect a tone. Prior to the fitting of ROCs the rating data were pooled by block (x 10) and across participants (x 12). However, at any one signal level the number of judgments provided by participants differed from each other, this likely a function of sensitivity. Values of *d′* and *d*_*a*_ as a function of signal level are displayed in Figure 8.

**Fig. 7.**
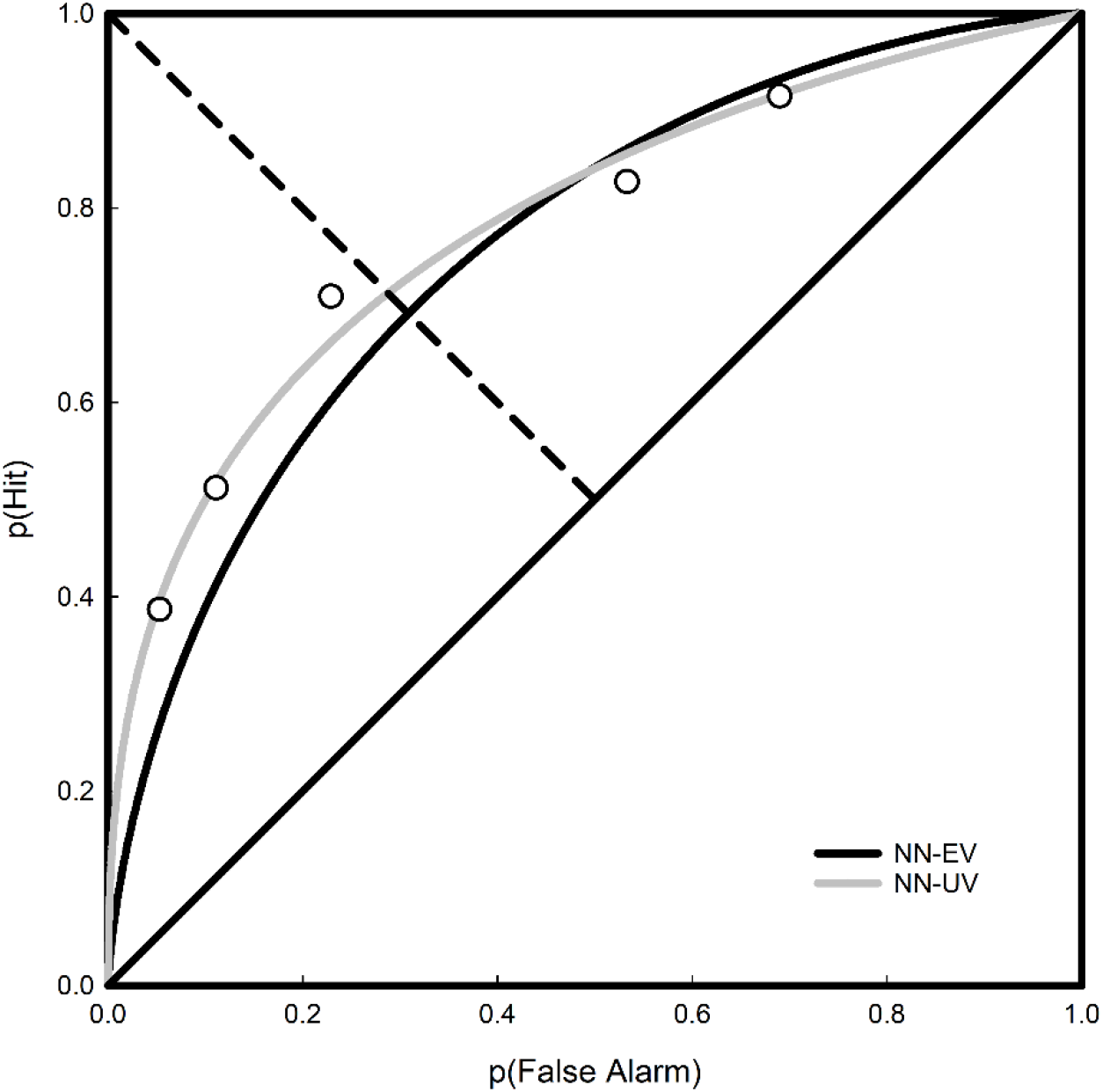
Receiver operating characteristic curves for pooled rating data at the 20 dB SPL signal level. The normal-normal equal-variance (black curve) and normal-normal unequal-variance (grey curve) models are plotted. Here, *d′* was estimated to be 1.186 and *d*_*a*_ to be 1.112.

**Fig. 8.**
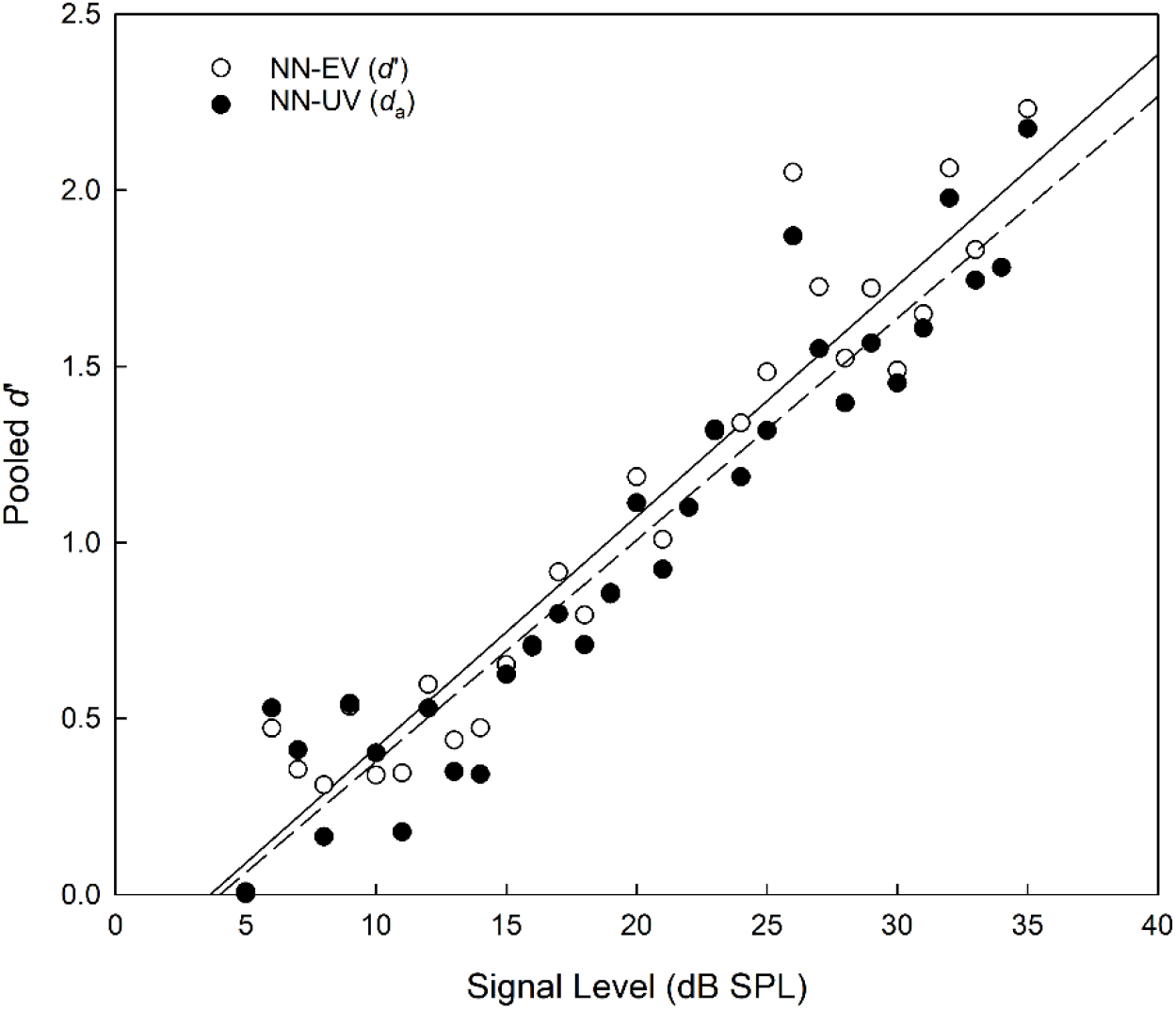
Pooled *d′* and *d*_*a*_ estimates as a function of signal level for two theoretical models, the NN-EV (open circles) and NN-UV (closed circles) models, respectively.

The ROCs displayed in Figure 7 are generated from pooled data collated across block and participant for the case when the signal level was 20 dB SPL, this being the closest to the group average of 19.75 dB SPL. With reference to Kaernbach’s SIAM procedure [2], when *t* = MRHR = .5, then p(FA) = .25 and p(Hit) = .75, yielding *d′* = z(.75) – z(.25) = 1.35 [9]. However, this only applies to the NN-EV model as it is only the NN-EV model for which *d’* is a relevant measure. For the NN-EV model, the MRHR occurs at the minor diagonal. This follows from the ROC curve being symmetrical. However, for the NN-UV model, this is no longer true, as the MRHR can veer off to one side at the location where the slope of the ROC curve is 1. Returning to Figure 8, the value of *d′* calculated for the NN-EV model is 1.19, which arguably is sufficiently close to a *d′* of 1.35. Goodness-of-fit indices for each signal level are displayed in Table 4, with chi-square (x^2^) values indicating poor fits as significant values indicate that model and data are not the same. Maximized log likelihood estimates were compared using the likelihood ratio (LR) test, allowing scrutiny of the extra parameter resident in the NN-UV model. Here, the extra parameter in the NN-UV model resulted in superior fits to the NN-EV model on approximately 2/3 of the data. Summing both the x^2^ and degrees of freedom values for each model (*re*: Table 4) and deriving the cumulative probability for each statistic again revealed that superiority of the NN-UV model, and that both the NN-EV and NN-UV models provided poor fits to the data (*p* < .001).

**Table 4.** Goodness of fit indices (x^2^) for the NN-EV and NN-UV models.

| Signal Level | Number of signals | NN-EV Model |  | NN-UV Model |  | NN-EV vs. NN-UV |  |
| --- | --- | --- | --- | --- | --- | --- | --- |
| | | $\chi^2$ | Log-Likelihood | $\chi^2$ | Log-Likelihood | Log-Likelihood ratio | $p$-value |
| 5 | 70 | 13.06** | -9372.44 | 3.94 | -9374.79 | 4.7 | 0.001 |
| 6 | 30 | 5.16 | -9307.19 | 3.12 | -9306.28 | 1.82 | 0.177 |
| 7 | 21 | 1.36 | -9293.26 | 0.11 | -9292.54 | 1.44 | 0.230 |
| 8 | 64 | 22.27*** | -9374.21 | 3.11 | -9365.21 | 18 | 0.001 |
| 9 | 44 | 5.32 | -9337.35 | 5.32 | -9337.35 | 0 | 0.999 |
| 10 | 56 | 7.63 | -9364.31 | 7.60 | -9364.31 | 0 | 0.999 |
| 11 | 105 | 32.53*** | -9447.05 | 9.32*** | -9435.06 | 23.98 | 0.001 |
| 12 | 95 | 9.67** | -9449.09 | 4.71 | -9446.47 | 5.24 | 0.022 |
| 13 | 96 | 12.36* | -9428.64 | 4.76 | -9424.79 | 7.7 | 0.005 |
| 14 | 244 | 67.95*** | -9705.69 | 10.19* | -9677.27 | 56.84 | 0.001 |
| 15 | 214 | 20.88*** | -9634.87 | 19.80*** | -9634.09 | 1.56 | 0.212 |
| 16 | 269 | 24.25*** | -9730.44 | 23.92 | -9730.4 | 0.08 | 0.777 |
| 17 | 397 | 75.32*** | -9956.08 | 20.02*** | -9929.94 | 52.28 | 0.001 |
| 18 | 354 | 20.94*** | -9894.02 | 26.35*** | -9888.6 | 10.84 | 0.001 |
| 19 | 326 | 33.36*** | -9836.28 | 33.22*** | -9836.25 | 0.06 | 0.807 |
| 20 | 457 | 50.26*** | -10025.66 | 25.86*** | -10013.53 | 24.26 | 0.001 |
| 21 | 378 | 69.96*** | -9930.37 | 45.51*** | -9916.02 | 28.7 | 0.001 |
| 22 | 283 | 22.74*** | -9743.09 | 22.74*** | -9743.09 | 0 | 0.999 |
| 23 | 360 | 58.66*** | -9816.43 | 19.49*** | -9800.44 | 31.98 | 0.001 |
| 24 | 248 | 73.97*** | -9663.64 | 26.42*** | -9644.46 | 38.36 | 0.001 |
| 25 | 152 | 45.48*** | -9494.15 | 11.01*** | -9481.04 | 26.22 | 0.001 |
| 26 | 178 | 60.29*** | -9452.36 | 5.14*** | -9439.88 | 24.96 | 0.001 |
| 27 | 112 | 32.48*** | -9409.24 | 5.35*** | -9399.92 | 18.64 | 0.001 |
| 28 | 98 | 20.10*** | -9397.24 | 8.39*** | -9390.56 | 13.36 | 0.001 |
| 29 | 124 | 26.97*** | -9417.56 | 7.10*** | -9407.12 | 20.88 | 0.001 |
| 30 | 88 | 3.97 | -9382.52 | 3.33 | -9382.11 | 0.82 | 0.365 |
| 31 | 78 | 9.94* | -9363.28 | 9.59* | -9362.83 | 0.9 | 0.343 |
| 32 | 99 | 9.67* | -9357.73 | 7.40 | -9355.78 | 3.9 | 0.048 |
| 33 | 51 | 7.23 | -9321.05 | 6.51 | -9319.9 | 2.3 | 0.129 |
| 34 | 39 | 9.24 | -9311.75 | 9.24* | -9311.75 | 0 | 0.999 |
| 35 | 79 | 20.65*** | -9327.87 | 5.23 | -9324.2 | 7.34 | 0.007 |
| Statistics | M = 171.3 | $\Sigma = 873.67$ | - | $\Sigma = 393.80$ | - | - | - |
| * $p < .05$ ** $p < .01$ *** $p < .001$ | | | | | | | |

In allowing full ROCs to be constructed, the SIAM-Rating task also affords the quantification of response bias. Yes-No tasks have traditionally used measures of bias based upon the likelihood ratio, such as *β*, though these have been found to be dependent upon sensitivity (i.e., *d′*). Instead, bias estimates based on the criterion, such as *c*, are recommended [18]:

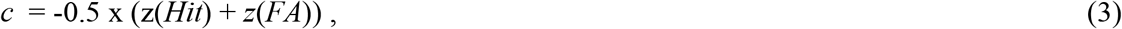

where negative values of *c* indicate a bias toward responding ‘yes’ (a liberal criterion), positive to ‘no’ (a conservative criterion), with *c* = 0 representing an unbiased observer. According to Equation 3, *c* is the average of the standardised Hit and FA probabilities, and represents the distance between the criterion and the no-bias point (i.e., *c* = 0), the latter being where the observer’s criterion is equidistant from the means of the signal distribution and the noise distribution. In the current study *c* was calculated for signal levels between 5 and 35 dB SPL using a correction prior to standardisation, such that *p*(Hit) = (Hit+0.5)/(signal trails + 1) and *p*(FA) = (FA+0.5)/(noise trails + 1) [18]. The mean value of *c* across the 31 signal levels was 0.035 (*SD* = 0.344, *Min* = 0.00, *Max* = -0.84), with a one-sample *t*-test indicating that the mean value of *c* was not significantly different from zero (*t*(30) = .793, *p* = 217).

### Psychometric Function

As for Figure 3 and the SIAM-TT task, a non-adaptive psychometric representation of the SIAM-YN data was generated using Equation 1 (*re*: Figure 9). Here, the area of each point of the empirical psychometric function reflects the number of trials used to calculate the point, with the mean number of trials across the points being 149.2 (*SD* = 148.1, *Min* = 2, *Max* = 509). The goodness-of-fit for Equation 1 was *R*^2^ = 0.98, while the mean goodness-of-fit across the 27 individual functions was *R*^2^ = 0.93 (*SD* = 0.08). With reference to the 50% point on the ordinate, a value of 19.06 dB SPL is obtained, within about 1 dB of the estimates reported in Table 3, and with no statistical significance between them (*p* > .05).

**Fig. 9.**
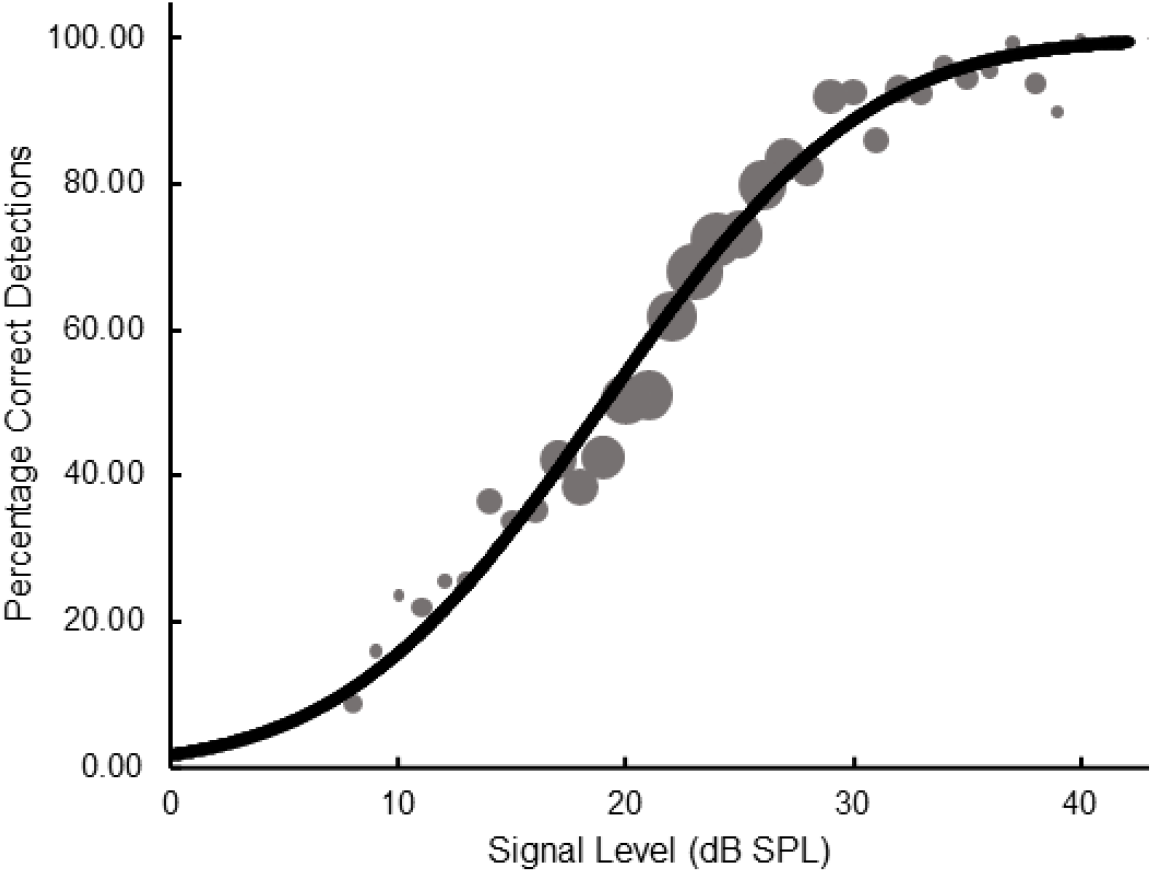
Psychometric function plotting percentage correct detections as a function of signal level for pooled data obtained from the SIAM-YN task. The slope parameter is equal to 0.83.

### Temporal Analysis

The mean number of trials and mean time (seconds) it took participants to get to 15 turnarounds are presented in Table 5. As for Experiment 1, the accuracy of these figures is slightly biased by measurement precision, though this should not affect one task any more than the others. The superscripted letters in Table 4 indicate, by scrutiny across a single row, significant differences across the three tasks. Of note, the 2-IFC task required a significantly greater number of trials and took longer to finish a block of trials than both the SIAM-YN and SIAM-Rating tasks. Considering seconds-per-trial, the SIAM-YN task was significantly faster than either the 2-IFC task or the SIAM-Rating task. The last row in Table 4 indicates that the SIAM tasks had significantly lower trials/turnaround ratios than the 2-IFC task.

**Table 5.** Group means indicating the number of trials, time taken, seconds-per-trial, and trials-per-turnaround, across the three tasks. Superscripted letters should be referenced across a single row, and indicate significant differences (*p* < .05) across tasks. Standard deviations are in parentheses.

|  | Task |  |  |
| --- | --- | --- | --- |
|  | a) 2-IFC | b) SIAM-YN | c) SIAM-Rating |
| Trials | 74.29<sup>b,c</sup> (2.98) | 40.02<sup>a</sup> (2.48) | 38.53<sup>b</sup> (3.53) |
| Block Time (secs) | 235.29<sup>b,c</sup> (14.97) | 110.16<sup>a</sup> (9.25) | 111.19<sup>b</sup> (8.41) |
| Secs/Trial (secs) | 3.17<sup>b,c</sup> (0.14) | 2.76<sup>a,c</sup> (0.15) | 2.90<sup>a,b</sup> (0.17) |
| Trials/Turn | 4.95<sup>b,c</sup> (0.20) | 2.67<sup>a</sup> (0.17) | 2.57<sup>a</sup> (0.23) |

## Discussion

The aims of Experiments 1 and 2 were to further validate the SIAM-YN task, and to assess two SIAM-YN modifications, the SIAM-TT and the SIAM-Rating tasks. All three versions of the SIAM task will be assessed with reference to Treutwein’s [7] three criteria for evaluating threshold estimation procedures. First, are the threshold estimates valid and reliable? Second, can the task be easily implemented? Third, given the context, is the task more efficient than the alternatives. Additionally, Shepherd et al. [10] added a fourth criterion to these three: is the procedure simple enough for participants to rapidly comprehend? These criteria will be kept to the fore when considering the current evaluation of the SIAM-YN task and its two modifications.

### Further evaluation of the SIAM-YN Task

Estimates of mean absolute threshold were not significantly different between the 2-IFC task and the SIAM-YN task. For Experiment 1 the mean difference between the two tasks was trivial (0.37 dB SPL, *re*: Table1), with the standard deviation approximately 20% lower for the SIAM-YN task. A similar pattern was reported in Experiment 2 (*re*: Table 3), with a difference of 0.23 dB SPL between the two tasks, though for this data the standard deviation is slightly lower for the 2-IFC task. In Experiment 2 the deployment of both ascending and descending trial sequences with the SIAM-YN task likewise reinforced the validity of the task through the construction of a psychometric function. Taken together, the analyses support the validity of the SIAM-YN threshold estimates and mirror the findings of Shepherd et al. [10] who reported mean thresholds of 22.7- and 22.98-dB SPL for the 2-IFC and SIAM-YN tasks respectively. Thus, in the psychoacoustics context there is further support of Kaernbach’s assertion that, when considering human data, the SIAM-YN task produces thresholds that are comparable to the 2-IFC task [2]. In terms of reliability, no statistical differences were found between the 2-IFC task thresholds measured across Experiments 1 and 2, nor those obtained with the SIAM-YN task, despite the potential for individual differences to affect the data.

### The SIAM-TT task

Threshold estimates for the SIAM-TT task were statistically indistinguishable from the 2-IFC and SIAM-YN tasks at the α = .05 criterion. Thus, if estimates from the 2-IFC task represent the benchmark, then arguably the SIAM-TT task has demonstrated convergent validity with the gold-standard procedure. Furthermore, the differences in standard deviations between the two tasks did not reach significance, with the SIAM-TT task’s estimate being 26% lower than the 2-IFC task. As a novel modification, there are no previously published studies reporting comparable data. The construction of a psychometric function using the raw data provided an additional test of convergent validity, this time with the one-interval yes-no task. Again, the estimate obtained using the SIAM-TT task was found to be in agreement.

Considering task efficiency, our data conclusively demonstrated that robust threshold estimates could be obtained using fewer trials than either its 2-IFC and SIAM-YN counterparts. Indeed, the SIAM-YN task required 45% more trials, and the 2-IFC task 53% more trials, than the SIAM-TT task. However, in terms of time per block, the SIAM TT task took approximately 13% longer than the SIAM-YN task, and its trials were over 40% longer. This is explained in part by having trials in which both tracks contained blanks, and therefore the participant had to wait the fixed time. Though the SIAM-TT task was on average approximately 20 seconds faster than the 2-IFC task, this difference was not statistically significant. Also of remark, unlike the 2-IFC procedures, which contain one bit of information [2] the SIAM-TT obtains two, and can thus be considered more efficient.

Finally, turning to participant-centred factors, our data showed no significant difference in learning effects across the three tasks, nor did the participants themselves report any difficulties with any of the three tasks, even though all participants were naïve. Part of this may come down to the use of laminated mats that clearly detailed both response and feedback regimens. In conclusion, the SIAM-TT task is an easily implemented psychoacoustical threshold estimation method that in terms of efficiency may possess significant advantages over the 2-IFC and SIAM-YN tasks. A further advantage is, if desired, the use of ascending and descending tracks to generate empirical psychometric functions to which theoretical models can be regressed.

### The SIAM-Rating task

As was found with the SIAM-TT task, the absolute threshold estimates obtained using the SIAM-Rating task were not statistically discernible from those calculated from the 2-IFC and SIAM-YN tasks. In particular, the pooled threshold estimate for the SIAM-rating task was not significantly different from the 2-IFC estimate, which was taken as a bench mark. Of further interest is the similarity between the SIAM-YN and Rating tasks, both being identical apart from an expanded number of response options for the latter. From the data it can be concluded that the change in response options does not impact threshold estimates, and so asks the question of the usefulness of the SIAM-Rating task over its parent task? Here, the obvious advantage is the ability of the SIAM-Rating task to generate a full ROC that is not based on a single point in ROC space, and can yield information on response bias. Pertinently, when the signal and noise distributions are both normally distributed and possess equal variances (i.e., the NN-EV model) then *d′* is assumed to be independent of response bias. While the binary response regime prevents these assumptions being tested in the SIAM-YN task, they can be tested in the SIAM-Rating task. Additionally, the SIAM-Rating task may better fulfil Kaernbach’s assumption [2] that response outcome is a reinforcer or punisher by varying the degree of reinforcement and punishment, an important area for future research [31].

Turning now to task efficiency, a block of the SIAM-Rating task took approximately the same number of trials as the SIAM-YN task, but was clearly more efficient than the 2-IFC task, needing approximately half as many trials, and consequently being completed in half the time. The SIAM-Rating task had, however, trials that were on average about 5% slower than the SIAM-YN task, but about 40% quicker than the two-interval trials of the 2-IFC task. Finally, and mirroring the SIAM-TT task, the SIAM-Rating task seemed as easy to learn as the other two tasks, as evidenced by the lack of a task x block interaction.

### Task Selection

For the psychophysicist, a central interest is sensory acuity and, furthermore, what accounts for limits in acuity. However, psychophysical techniques have been applied far beyond the primary interests of psychophysics, where efficiency may serve as the primary motivation to use adaptive procedures in the first place. It also needs to be acknowledged that when selecting tasks a number of modifications can be considered – the determination of signal level as a function of trial, when to halt a block of trials, and how to calculate threshold. However, as these customisations apply to all the tasks considered here, such modifications do not constitute selection criteria when choosing across these tasks. Rather, accuracy and efficiency are best used to inform choices of test, though the decision is complicated given the complimentary relationship between the two. Referencing the concept of ‘work’ in physics, Taylor and Creelman [21] proposed the ‘sweat factor’ metric, where trial number is multiplied by the variance of the threshold estimate. This metric applied to the current dataset shows the 2-IFC to be more exerting than the three SIAM tasks.

While the 2-IFC transformed up-down procedure has dominated adaptive testing over the last fifty years, the conclusion taken from the current data and that of Kaernbach’s [2] is that the SIAM-YN task is more efficient and is equally precise. In his advocacy of single-interval procedures, Kaernbach [2] goes as far to declare “…*it is superfluous to present more than one interval per trial*.” (p. 2653), though the degree to which this comment would hold for the SIAM-TT task presented here is uncertain. In terms of implementation, the 2-IFC transformed up-down procedure and all three of the SIAM procedures evaluated in the current study were easy to set-up, with either approach able to adjust stimulus levels by pen-and-paper if computer control were not available. At the same time, both approaches have strong theoretical backbones, as formulated by Levitt and Kaernbach [1,2]. This ease of implementation is a point that we will return to later.

Other adaptive procedures beyond those employed in this study exist, including maximum-likelihood methods (ML: e.g., ML YN task / QUEST and derivatives), parameter estimation by sequential testing (PEST), and non-parametric (e.g., Up-Down Transformed-Response) tasks. Computer simulations have consistently demonstrated the superiority of the MLE and PEST tasks over orthodox staircase methods, however, the same finding has not been reliably reported when human data is collected [13,15,17]. King-Smith et al. [22] suggest that the contrasting finding between human data and simulations could possibly be due to the greater volume of data that can be obtained with the latter, or because simulation assumptions (e.g., statistical independence of trials and invariant threshold) have been shown to be frail for human participants [23]. Further, the corrections required by MLE techniques such as QUEST to account for human factors such as attentional or memory lapses, and shifts in threshold, substantially increase the number of trials at the cost of efficiency [18]. As the ecological validity of simulated data is therefore open to question, and given the criticism of some MLE tasks [24,25], it may be useful to heed the concluding remarks of Kollmeier et al. [17]:

*However, because the observed differences in efficiency are relatively small and inconsistent, other experimental design criteria should be weighed more heavily than the efficiency of threshold estimation*. (p. 1861).

For example, the QUEST procedure necessitates the psychometric function to be described *a priori* and is vulnerable to parameter mismatches [8], and is thus best suited to cases in which previous results are available [26]. The simplicity of the staircase procedures and the SIAM-YN tasks is such that they can be administered without the use of a computer, nor are dependent on previous data.

### Strengths and Limitations

While the proposed modifications to the SIAM-YN task appear promising in light of future test development, the current findings must be interpreted with reference to the method and analytical approaches. First, in terms of the participants, the sample consisted of young university students who were motivated to learn, and while they were asked verbally if they had experienced noise-induced hearing loss in the past, this was not measured objectively using the audiogram. A strength of the current approach is the reliance upon human participants, as opposed to more commonly encountered computer-generated simulation data. While Kaernbach [2] provided the outputs of simulations when assessing the SIAM-YN task, the threshold estimates derived from Monte Carlo techniques are based on *a priori* determined probability distributions and parameters which represent both sensory and response variability, the latter of which can be problematic [27]. Simulations typically rely upon the conceptualisation of the ‘ideal’ observer, in which optimal performance is converged upon within the defined sensory contexts. While simulation has advantages, for example, being more easily conducted than human studies as-well-as providing important baseline data, the burden of modelling sensory and decision processes is a disadvantage that is not shared by using human observers [28].

An additional caveat is our use of the 3-down 1-up 2-IFC task rather than the more common 2-down 1-up task, with the former requiring a lower number of trials for a fixed number of turn-arounds [29]. Our selection of the 3-down 1-up regime was undertaken to bring the convergence probabilities of the SIAM-YN (*p* ≈ 0.83) and 2-IFC (*p* ≈ 0.80) tasks closer together, so they would be closer matched in terms of accuracy. As such, using the 2-down 1-up (*p* ≈ 0.71) rule would result in a skewed comparison as its threshold estimates would be less accurate than those calculated from the SIAM tasks. Indeed, Green [30] eschewed the 2-down 1-up regime in favour of regimes that track higher performance criteria which, he argued, are both more accurate and efficient. As detection is a probabilistic process any estimate of absolute threshold will be variable, with the variability proportional to target performance [18]. The performance ‘sweet point’ proposed by Green [30] to minimise statistical bias is at the 91% correct point on the Yes-No psychometric function, a full 20% higher than the 2-down 1-up regime’s 71% target. Thus, the adoption of the 3-down 1-up regime over the 2-down 1-up regime permits gains in both precision and efficiency, as demonstrated in the auditory context by Kollmeier et al. [17].

## Conclusion and Future Directions

In these two studies we present further evidence that the SIAM-YN procedure can produce reliable and valid estimates of absolute threshold. While the 2-IFC staircase task is ubiquitous in sensory and perceptual research, its application reaches far beyond psychophysics. In these other arenas the estimation of thresholds may be secondary to other measurements, and therefore efficiency may be a key consideration when deciding which task to employ. Based on the current and previous [2,11] research, we would argue that the SIAM-YN task is a worthy replacement for the 2-IFC task for the estimation of absolute thresholds. Considering the two modifications proposed here, the SIAM-TT task would be of utility when both descending and ascending tracks are desired, or for when psychometric functions are desirable. Further, the SIAM-TT task could be utilised with twin-track SIAM-Rating tasks. The SIAM-Rating task, with its extended number of response options and ability to generate a full ROC without decreases in efficiency, may be considered an able replacement for the standard SIAM-YN task itself. Future experimentation of the SIAM tasks and their starting parameters (e.g., step-size, staircase type, starting level) would be useful, along with comparisons to tasks utilising Bayesian staircases.

### Supporting information

SDT Assistant download:

https://hautus.org/sdt-assistant.php

## Acknowledgements

We would like to thank Miriam Stocks and Maraya Brogli for assisting with data collection and collation.

## Author Contributions

Conceptualization: Daniel Shepherd, Michael Hautus.

Formal analysis: Daniel Shepherd, Michael Hautus.

Investigation: Daniel Shepherd, Michael Hautus.

Methodology: Daniel Shepherd, Michael Hautus.

Project administration: Daniel Shepherd, Michael Hautus.

Software: Michael Hautus.

Visualization: Daniel Shepherd, Michael Hautus.

Writing – original draft: Daniel Shepherd, Michael Hautus.

Writing – review & editing: Daniel Shepherd, Michael Hautus.

